# Flow-induced bending of flagella controls bacterial surface behavior

**DOI:** 10.1101/2025.01.07.631359

**Authors:** Jessica-Jae S. Palalay, Joseph E. Sanfilippo

## Abstract

Bacteria engage in surface-specific behaviors that are assumed to be driven by biological signaling. However, surface behaviors could be controlled by mechanical reorientation of bacterial appendages. Here, we use microfluidics and flagellar labeling to discover how shear force bends flagella to control surface behavior of the human pathogen *Pseudomonas aeruginosa*. By imaging flagellar rotation and using mutants with paralyzed flagella, we establish that flagellar rotation promotes surface departure in host-relevant shear regimes. Our single-cell experiments reveal two distinct subpopulations in flow: cells with their flagellum positioned upstream and cells with their flagellum positioned downstream. Shear force bends upstream flagella around the cell and blocks rotation. In contrast, downstream flagella can continue to rotate after surface arrival. Cells with downstream flagella depart the surface more frequently than cells with upstream flagella, indicating how flow direction can determine bacterial cell fate on surfaces. Together, our results demonstrate how the geometric relationship between flow and cell appendages can generate subpopulations and control surface behaviors.

## Introduction

Mechanical features of the environment impact bacterial behavior (1–3). For example, pathogenic bacteria are influenced by shear forces and surface association when infecting heart valves or catheters (4–6). Recent discoveries have led to the emerging paradigm that bacteria can respond to environmental mechanics through active sensing and signaling (2, 7–10). As the cell wall makes bacterial cells rigid, researchers hypothesized that bacterial mechanosensing is mediated by flexible appendages such as flagella and type IV pili (7, 11–16). While there is evidence that flagella and type IV pili control mechanical responses (17, 18), the mechanisms that underly these responses remain controversial and unclear (19, 20). Considering the importance of mechanical host-pathogen interactions, there is an urgent need to better understand how mechanical forces modulate bacterial behaviors.

Conceptually, there are at least three ways that environmental mechanics could impact bacterial behavior (2, 3). First, a mechanical stimulus could be sensed by a cell surface component, which could lead to changes in internal signaling. There is evidence that flagella and type IV pili use this mechanism to sense surface contact and trigger surface associated behaviors (14, 16–18). Second, a mechanical stimulus could impact the local concentration of small molecules, which could be sensed by the cell and trigger internal signaling. There is evidence that shear rate impacts behavior by replenishing or washing away small molecules such as autoinducers or hydrogen peroxide, which ultimately are sensed by the cell (9, 21–23). Third, a mechanical stimulus could induce a conformational change in the cell or a cell surface component that subsequently impacts behavior without altering internal signaling. There is evidence that shear force modulates adhesion by tipping cells over or stretching a cell surface adhesin that displays catch bond properties (24–26). As there is evidence for many forms of bacterial mechanoregulation, it is important to consider various possibilities when characterizing a bacterial response.

During infection, bacteria must contend with host-generated fluid flow (4, 5, 27). Using microfluidic systems, recent studies demonstrated how flow impacts antimicrobial resistance (28), quorum sensing (22, 29), gene expression (9, 14, 21, 23), and surface motility (25, 26, 30). Perhaps the most obvious impact of flow is on adhesion, where the shear force associated with flow can remove cells from a surface. However, the study of adhesion in flow revealed the counterintuitive result that increasing flow often enhances adhesion (25, 26, 31, 32). In lower shear regimes, *Pseudomonas aeruginosa* cells use type IV pilus retraction to promote surface departure by tilting themselves away from the surface (26). Higher shear forces overcome pilus-dependent tilting, push *P. aeruginosa* cells closer to the surface, and increase their surface residence time (26). This example highlights how the interaction between cell-generated forces and environmental forces can lead to unexpected outcomes.

To study the interaction between cell-generated forces and environmental forces, we examined how *P. aeruginosa* adhesion is impacted by flagellar rotation and flow. Using mutants with paralyzed flagella, we show that flagellar rotation promotes both surface arrival and departure in host-relevant shear regimes. Combining microfluidics and flagellar labeling, we demonstrate that flow can bend the flagellum and block rotation. By tracking single cells in flow, we discover two distinct subpopulations: cells with their flagellum positioned upstream and cells with their flagellum positioned downstream. By independently modulating flow intensity and solution viscosity, we establish that shear force bends upstream flagella and halts flagellar rotation. In contrast, downstream flagella are not bent by shear force and can continue to rotate after surface arrival. Importantly, cells with downstream flagella leave the surface more often than cells with upstream flagella. Collectively, our results reveal how the direction of environmental shear force interacts with the force of flagellar rotation to control bacterial surface behavior.

## Results

To investigate how the flagellum impacts initial *P. aeruginosa* surface interactions, we generated mutant strains lacking functional flagella (Figure 1A). To confirm our strains, we measured swimming motility with a traditional swim plate assay (Figure S1). While wildtype (WT) cells were motile, Δ*fliC* mutant cells lacking the flagellar filament exhibited no motility (Figure S1). Additionally, mutants lacking either flagellar stator complex (MotAB or MotCD) had decreased motility (Figure S1), confirming that both stator complexes contribute to flagellar rotation (33–36). Consistent with previous reports (33–36), our mutant lacking both *motAB* and *motCD* exhibited no swimming motility (Figure S1). Thus, we chose to use Δ*fliC* to examine the role of the flagellum and Δ*motAB* Δ*motCD* to examine the role of flagellar rotation.

**Figure 1:**
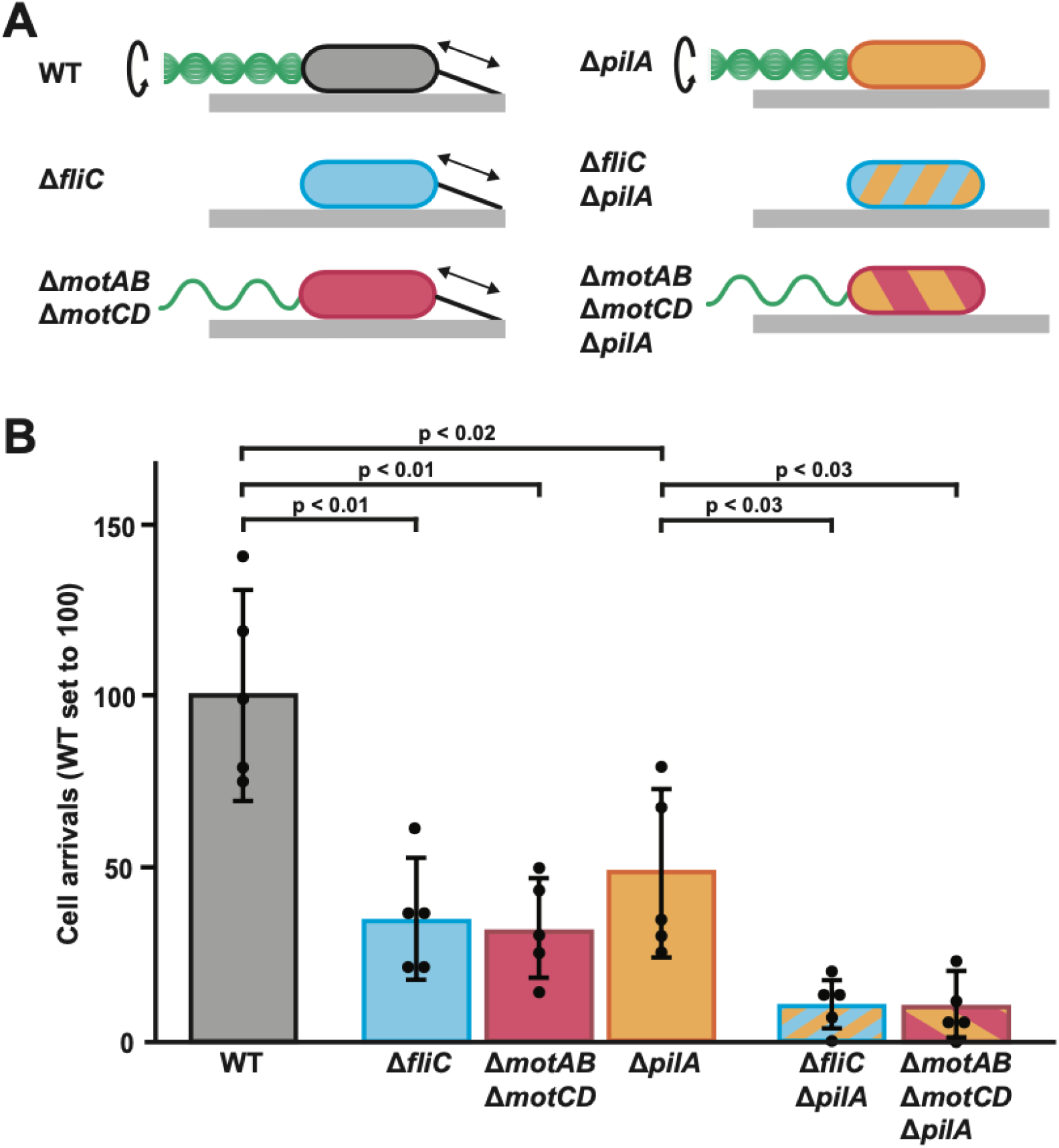
Flagellar rotation promotes *P. aeruginosa* surface arrival in flow. (A) Representation of wildtype (WT) and mutant strains used in this study. Cells either have a rotating flagellum (top row), lack a flagellum (middle row), or have a non-rotating flagellum (bottom row). Flagella in the diagram are located on the left side of the cells and are in green. Cells also have dynamic type IV pili or lack type IV pili. Pili in the diagram are located on the right side of the cells and are in black. (B) Quantification of cell arrivals for WT and mutant strains flowed into microfluidic devices at a shear rate of 800 s^-1^. Cell arrivals were determined by how many cells landed on the surface and remained attached for at least 2 seconds. Measurements were taken over 30 seconds and WT cell arrivals were normalized to 100. Quantification shows the average and standard deviation of five biological replicates. P-values were calculated with the Student’s T-test.

To determine if flagellar rotation impacts surface arrival in flow, we loaded cells into a syringe at a mid-log concentration and introduced them into empty microfluidic channels at a shear rate of 800 s^−1^. Surface arrival was recorded when a cell adhered to the surface and remained for at least 2 seconds (Figure 1B, S2). Compared to WT, Δ*fliC* cells had approximately 65% less arrivals and Δ*motAB* Δ*motCD* cells had approximately 70% less arrivals (Figure 1B). Thus, flagellar rotation (and not simply the presence of a flagellum) has a major role in *P. aeruginosa* surface arrival in flow. Type IV pili also have an important role in surface arrival (26). We confirmed this result using a Δ*pilA* mutant (which lacks the pilus filament) (Figure 1B) and generated combinations of mutations to test the interaction between type IV pili and flagellar rotation (Figure 1A). Compared to Δ*pilA* cells, Δ*fliC* Δ*pilA* had approximately 80% less arrivals and Δ*motAB* Δ*motCD* Δ*pilA* cells had approximately 80% less arrivals (Figure 1B). Together, these results establish that flagellar rotation acts independently of type IV pili to promote *P. aeruginosa* surface arrival in flow.

Does flagellar rotation impact surface departure in flow? As a rotating flagellum generates mechanical force, we hypothesized that flagellar rotation has a role in surface departure. To quantify surface departure, we measured the surface residence time (the time between surface arrival and departure) of cells exposed to a shear rate of 800 s^−1^. While WT cells had a short residence time (median = 2 min), Δ*fliC* cells (median = 4.8 min) and Δ*motAB* Δ*motCD* cells (median = 4.2 min) exhibited statistically longer residence times (Figure 2). Based on these results, we hypothesize that flagellar rotation has an important role in *P. aeruginosa* surface departure in flowing environments.

**Figure 2:**
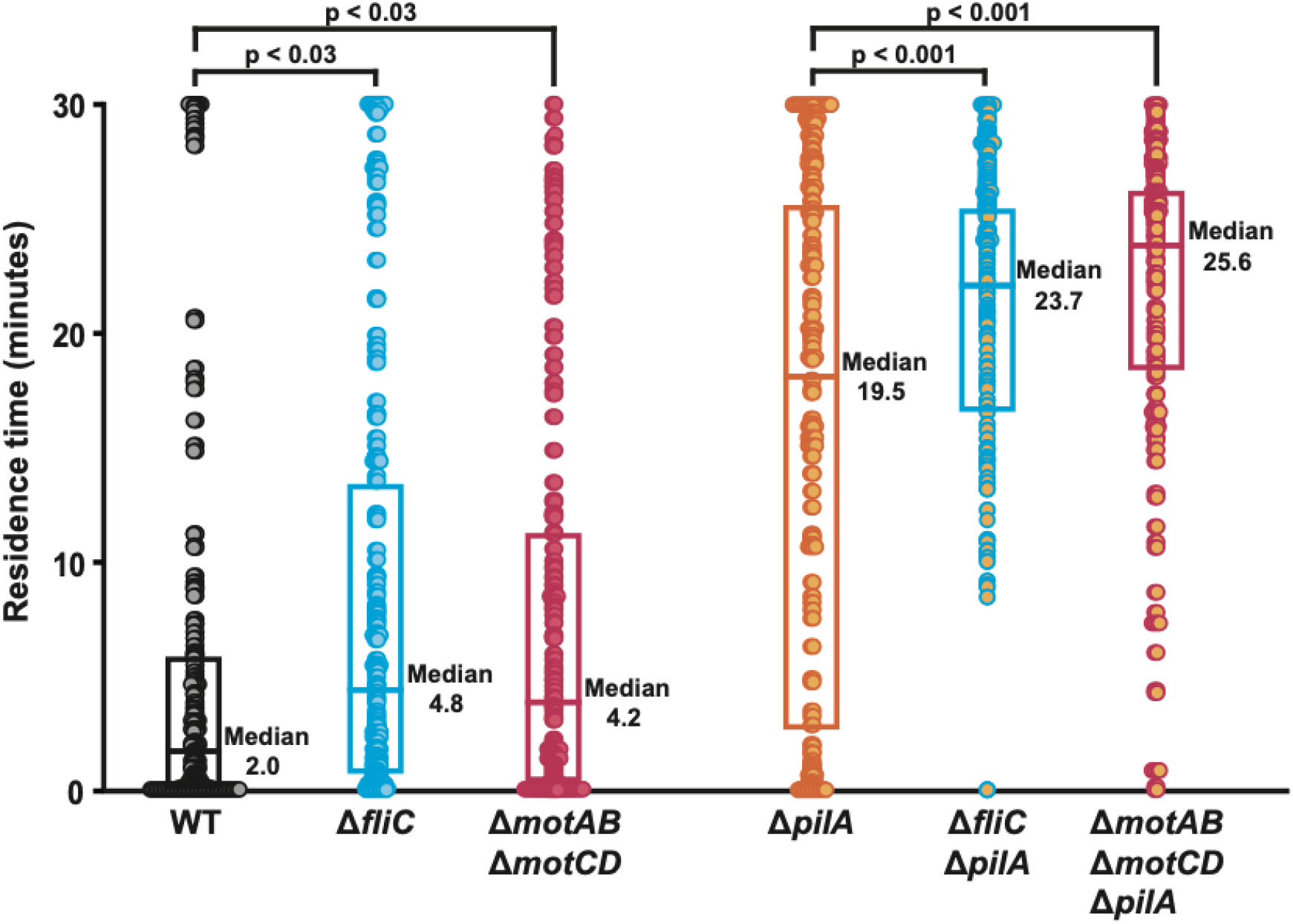
Flagellar rotation promotes *P. aeruginosa* surface departure in flow. Surface residence time of WT and mutant cells flowed into microfluidic devices at a shear rate of 800 s^-1^. Residence time is the time between surface arrival and departure. Each data point represents the residence time of one cell. Three biological replicates were performed and 150 cells (50 from each replicate) of each bacterial strain were chosen at random for quantification. P-values were calculated with the Student’s T-test, and the boxplot represents the 25^th^ percentile, median, and 75^th^ percentile for each strain.

Our previous work established that type IV pili have an important role in surface departure (26). Confirming the role of type IV pili, Δ*pilA* cells (median = 19.5 min) had a much longer residence time than WT (Figure 2). Using our combination mutants, we tested how type IV pili and flagellar rotation interact during surface departure. Compared to Δ*pilA*, Δ*fliC* Δ*pilA* cells (median = 23.7 min) and Δ*motAB* Δ*motCD* Δ*pilA* cells (median = 25.6 min) exhibited statistically longer residence times (Figure 2). Thus, we conclude that type IV pili and flagellar rotation contribute independently to surface departure. When focusing on the first 2 minutes, we noticed that while 24% of Δ*pilA* cells departed, only 1% of Δ*fliC* Δ*pilA* cells and 2% of Δ*motAB* Δ*motCD* Δ*pilA* cells departed (Figure S3). These experiments support the hypothesis that flagellar rotation promotes cell departure during early surface interactions in flow.

Do surface-attached cells rotate their flagella? To visualize flagellar rotation of recently adhered *P. aeruginosa* cells, we introduced a cysteine point mutation (T394C) in the flagellar filament protein FliC (37, 38). Then, we used a thiol-reactive Alexa488 maleimide dye to fluorescently label the flagellum (37). We observed that the majority of cells (∼70%) had 1 flagellum (Figure S4) with a median length of 3.4 µm (Figure S5). To examine flagellar rotation immediately after surface arrival, we introduced cells into a microfluidic device at a shear rate of 800 s^-1^ while simultaneously imaging fluorescently labeled flagella. As flagellar rotation is much faster (39–41) than the exposure time we used, we reasoned that rotating flagella would appear as rectangular-shaped blurs (Figure 3A). In contrast, non-rotating flagella appear in two dimensions as waveform shapes (Figure 3A). We observed that approximately 50% of WT flagella were not rotating 1 second after cell surface arrival (Figure 3B). In contrast, the other 50% of WT flagella rotated for many seconds before stopping (Figure 3B). Confirming the role of MotAB and MotCD in flagellar rotation, 100% of flagella from Δ*motAB* Δ*motCD* cells did not rotate (Figure 3B). Our results establish that there are two types of recently adhered *P. aeruginosa* cells: those with rotating flagella and those with non-rotating flagella.

**Figure 3:**
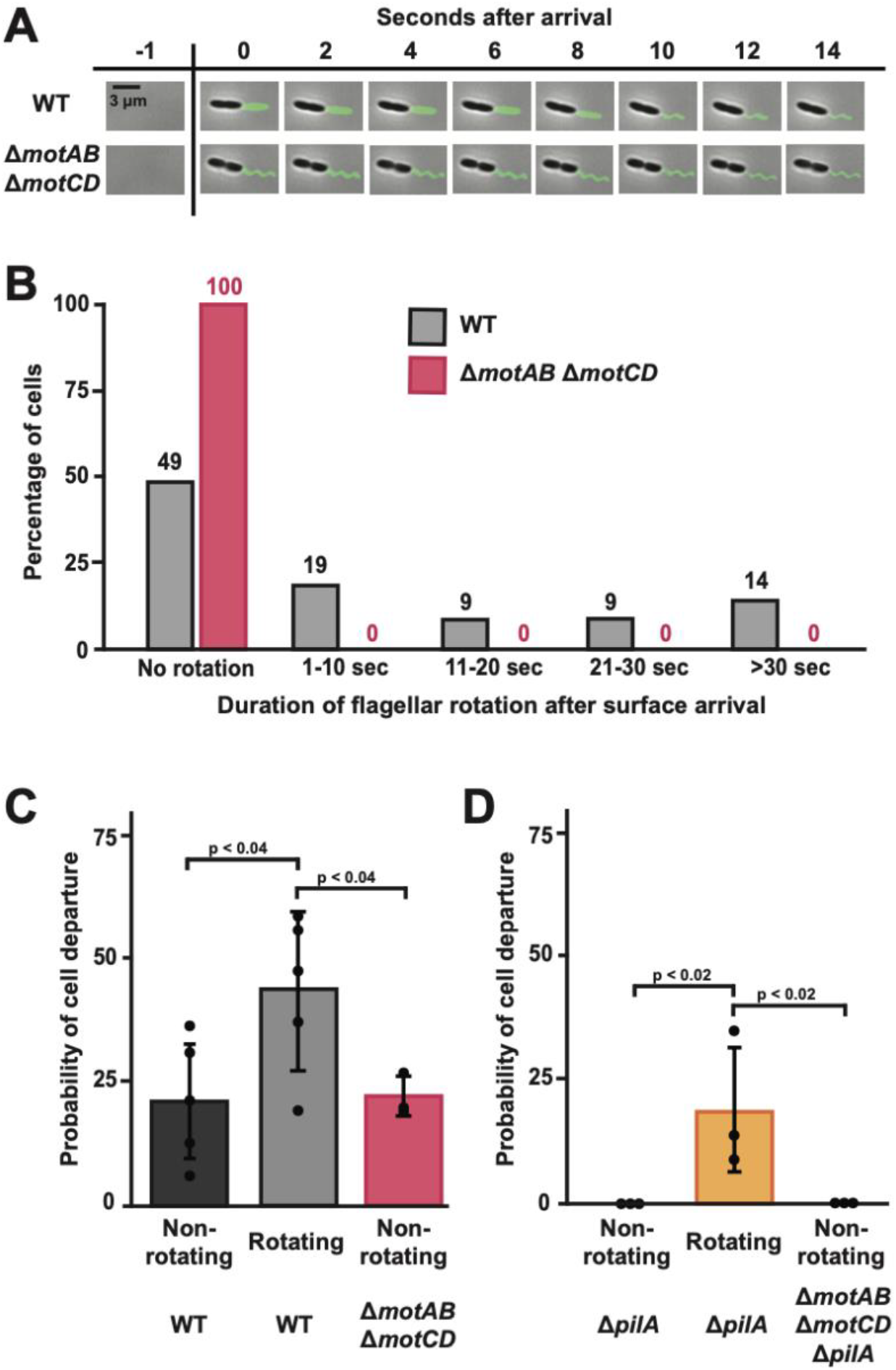
Cells with rotating flagella are more likely to depart the surface. (A) Representative images of surface-attached WT (top) and Δ*motAB* Δ*motCD* (bottom) cells with fluorescently labeled flagella. WT cell has a rotating flagellum, while Δ*motAB* Δ*motCD* cell does not have a rotating flagellum. Scale bar, 3 µm. (B) Duration of flagellar rotation after surface arrival of WT and Δ*motAB* Δ*motCD* cells. Five biological replicates were performed and a total of 550 WT cells and 125 Δ*motAB* Δ*motCD* cells were chosen for quantification and representation. Mean flagellar rotation of WT and Δ*motAB* Δ*motCD* cells are statistically different with *P* < 0.001. (C) Probability of surface departure of WT and Δ*motAB* Δ*motCD* cells. (D) Probability of surface departure of Δ*pilA* (orange line and shading) and Δ*motAB* Δ*motCD* Δ*pilA* (red line and orange & red shading) cells. All experiments were performed at a shear rate of 800 s^-1^. Cells were categorized as having rotating flagellum or non-rotating flagellum and the percentage of cell departures was calculated. Quantification shows the average and standard deviations of at least three biological replicates. P-values were calculated with the Student’s T-test.

Do cells with rotating flagella have a higher frequency of surface departure? To examine the relationship between flagellar rotation and surface departure, we quantified the probability of surface departure of WT cells with rotating flagella and non-rotating flagella. We observed that WT cells with rotating flagella had a 45% chance of surface departure, while WT cells with non-rotating flagella only had a 20% chance (Figure 3C). Consistent with the hypothesis that flagellar rotation promotes departure, Δ*motAB* Δ*motCD* cells (which lack flagellar rotation) had only a 21% chance of surface departure (Figure 3C). We reasoned that the residual surface departure in cells with non-rotating flagella was due to the effects of type IV pili. In support of our hypothesis, 0% of Δ*pilA* cells with non-rotating flagella departed the surface, while 19% of Δ*pilA* cells with rotating flagella departed the surface (Figure 3D). Similarly, 0% of Δ*motAB* Δ*motCD* Δ*pilA* cells (which lack flagellar rotation and type IV pili) departed the surface (Figure 3D). Together, our results demonstrate that flagellar rotation promotes cell surface departure in flowing environments.

While imaging labeled flagella, we observed two distinct subpopulations of cells: cells with their flagella facing upstream and cells with their flagella facing downstream (Figure 4A). As approximately 50% of cells had upstream flagella and 50% of cells had downstream flagella (Figure S6), we had a unique opportunity to examine how the directionality of flow impacts flagellar rotation. Flagella facing downstream rotated for a median of 16 seconds after cell surface arrival before stopping (Figure 4B). In contrast, all flagella facing upstream were not rotating in the first frame after cell surface arrival (Figure 4B). The sole exception was a cell that was shielded from flow by another tilted cell which was located directly in front of it (Figure S7).

**Figure 4:**
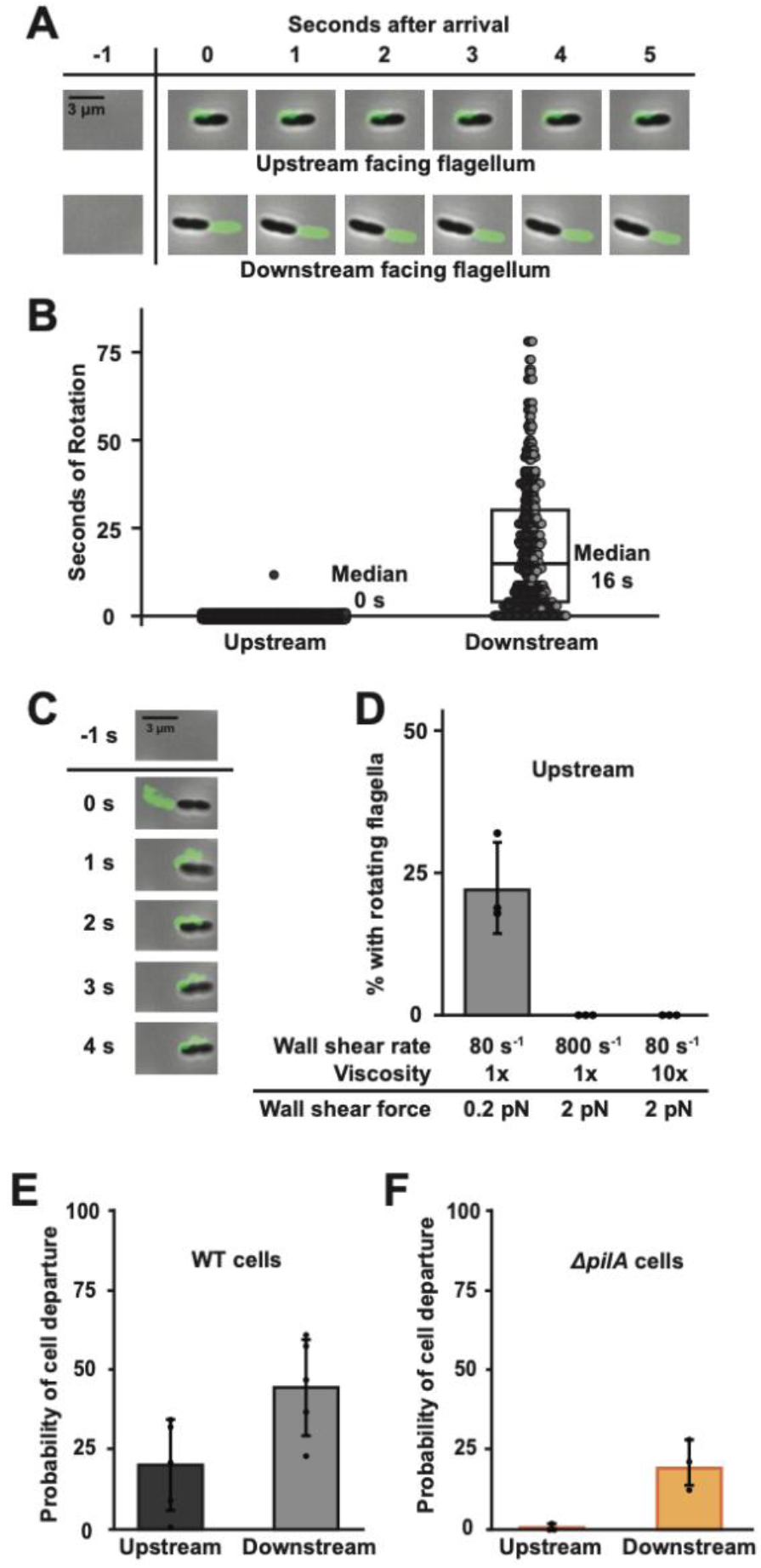
Shear force bends upstream facing flagella and blocks their rotation. (A) Representative images of surface-attached WT cells with an upstream facing flagellum or a downstream facing flagellum. Scale bar, 3 µm. (B) Duration of flagellar rotation of WT cells. Each data point represents one flagellum. Five biological replicates were performed and 528 WT cells were chosen for quantification. Boxplot represents the 25^th^ percentile, median, and 75^th^ percentile. (C) Images of a cell as flow bends its flagellum. Scale bar, 3 µm. (D) Percentage of cells with upstream flagella that are rotating during exposure to different shear forces. Shear force was increased either by changing shear rate or fluid viscosity. Shear rate was modified by changing the flow rate of the syringe pump. 10x viscosity was generated by adding 15% Ficoll, which has been shown previously to modify local viscosity (9, 42). (E) Probability of cell departure of WT cells with upstream facing or downstream facing flagella. Mean cell departures were statistically different with *P* < 0.04. (F) Probability of cell departure of Δ*pilA* cells with upstream facing or downstream facing flagella. Mean cell departures were statistically different with *P* < 0.05. Quantification shows the average and SD of at least three biological replicates. Shear rate was 800 s^-1^, except where noted. P-values were calculated with the Student’s T-test.

Based on these results, we hypothesize that flow rapidly blocks rotation of upstream facing flagella.

How does flow block rotation of upstream facing flagella? During our quantification of flagellar rotation time, we noticed that upstream flagella are bent around the cell body (Figure 4A, S8). After closer examination, we observed that upstream flagella typically bend around the cell body in the first second after surface arrival (Figure 4C). We hypothesized that flagellar bending was due to the shear force associated with flow. As shear force depends on shear rate and solution viscosity (9), we tested our hypothesis by independently modulating both variables. As increasing shear rate or viscosity increased the frequency of bent flagella, we concluded that flagellar bending was due to shear force (S8). To test if shear force is responsible for blocking rotation of upstream flagella, we quantified rotation while independently modulating shear rate and viscosity. While 23% of upstream flagella rotated at a shear rate of 80 s^-1^, no upstream flagella rotated when the shear rate was increased to 800 s^-1^ or the viscosity was increased by adding Ficoll (Figure 4D). Thus, upstream flagella are bent around the cell body and prevented from rotating by the shear force associated with flow.

Here, we made two important discoveries: flagellar rotation promotes surface departure and shear force blocks rotation of upstream flagella. Combining these two ideas, we hypothesized that cells with upstream flagella would be less likely to depart the surface in flow. In support of our hypothesis, WT cells with upstream flagella were less likely (20%) to depart the surface than cells with downstream flagella (45%) (Figure 4E). As type IV pili also promote surface departure, we repeated this experiment with Δ*pilA* cells. Similarly, Δ*pilA* cells with upstream flagella were less likely (1%) to depart the surface than cells with downstream flagella (19%) (Figure 4F). Based on this data, we conclude that cells with their flagella facing upstream are less likely to depart the surface than cells with their flagella facing downstream. Together, our findings establish that shear force bends flagella, blocks rotation, and ultimately controls surface behavior of *P. aeruginosa*.

## Discussion

Our experiments demonstrate how flow and flagellar rotation interact to control surface departure of *P. aeruginosa*. Using mutants with paralyzed flagella, we showed that flagellar rotation drives surface arrival in flow (Figure 1). While tracking cells after surface arrival, we learned that flagellar rotation also promotes surface departure in flow (Figure 2). By combining microfluidics and flagellar labeling, we discovered that flagella can rotate after surface arrival and cells with rotating flagella are more likely to depart the surface (Figure 3). Additionally, we discovered that shear force bends upstream facing flagella and blocks their rotation (Figure 4). Together, these experiments lead us to a model where cells with downstream facing flagella are more likely to depart the surface than cells with upstream facing flagella (Figure 5). Our results highlight how environmental and cell-generated mechanics can interact to yield unexpected surface behaviors.

**Figure 5:**
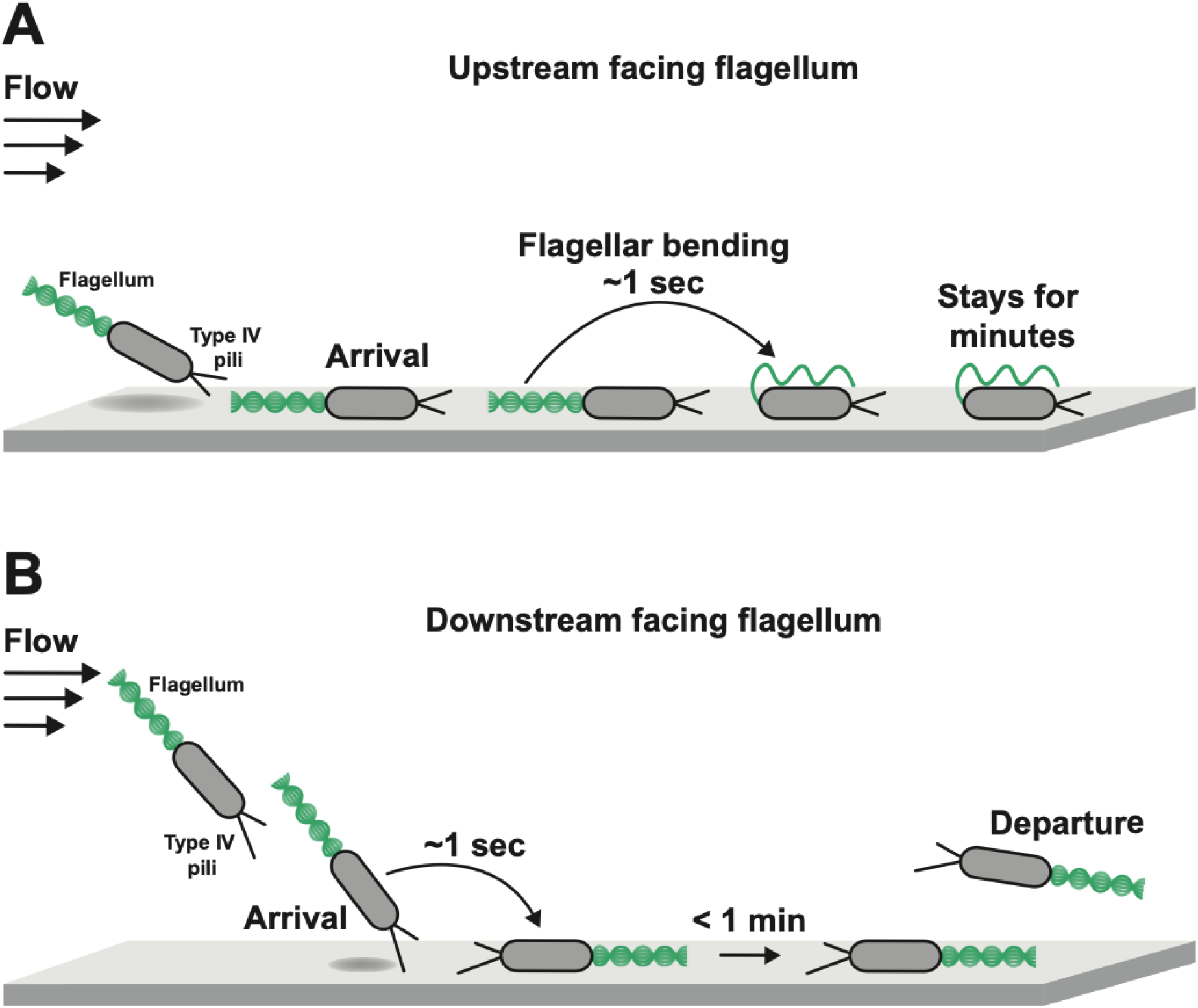
Flow-induced bending of flagella controls bacterial surface behavior. *P. aeruginosa* cells that arrive to the surface in flow can either have upstream facing (A) or downstream facing (B) flagella. (A) Cells can arrive on the surface with their flagella facing upstream into the flow. Quickly after arrival, the shear force associated with flow bends their flagella around their cell body and blocks rotation. As a consequence, these cells typically remain adhered to the surface. (B) Alternatively, cells can tip over as they arrive on the surface and have their flagella facing downstream. The flagella of these cells can continue to rotate and these cells are likely to quickly depart the surface.

How does shear force block flagellar rotation? By independently changing flow intensity and solution viscosity, we discovered that increasing shear force blocks rotation of upstream facing flagella (Figure 4D). Additionally, we discovered that increasing shear force bends upstream facing flagella (Figure S8B). These results lead us to the hypothesis that flagellar bending traps the flagellar filament next to the cell, which physically prevents rotation. An alternative model where the cell “jump ropes” over a rotating flagellum is possible (43, 44), but we did not observe this behavior in our experiments. As we did not observe motion of upstream flagella after a few seconds on the surface, we conclude that rotation had truly stopped. While we could not determine if the rotation of upstream flagella was stalled or permanently stopped (45), we did not observe any events where upstream flagella rotated again after stopping. Thus, our data support a model where shear force bends upstream flagella, traps the filament between the surface and the cell, and ultimately blocks rotation.

How do environmental mechanics impact bacterial behavior? The data presented here support the conclusion that environmental shear force can directly impact behavior without the need for intracellular signaling. In our model, physical bending blocks flagellar rotation, which controls whether or not a cell departs the surface. In contrast, there is evidence that other forms of mechanoregulation require intracellular signaling. For example, mechanosensing of surface contact by type IV pili and flagella controls intracellular signaling that triggers synthesis of a surface adhesin (17, 18). Additionally, flow-driven transport of hydrogen peroxide overcomes cell scavenging to trigger an intracellular signaling pathway (21, 23). When comparing and contrasting flagellar bending, mechanosensing, and flow-driven transport, it becomes clear that environmental mechanics can impact bacterial behavior in ways that appear phenotypically similar but mechanistically different. Moving forward, researchers should continue to search for new forms of bacterial mechanoregulation while understanding that the mechanisms underlying these processes may not require active sensing and signaling.

## Acknowledgements

We thank Sampriti Mukherjee for her gift of the *P. aeruginosa* T394C strain for flagellar labeling. We thank Anu Sharma, Piyush Sharma, Alex Shuppara, Gilberto Padron, Evan Johnson, Sizhe Chen, and Dan Kearns for helpful discussions and comments on the manuscript. This work was supported by grants K22AI151263 and R35GM155443 from the National Institutes of Health to J.E.S.

## Contributions

J.-J.S.P. and J.E.S. designed research. J.-J.S.P. performed research. J.-J.S.P. and J.E.S. analyzed data. J.-J.S.P. and J.E.S. wrote the paper.

