## Supplemental Information for "Flow-induced bending of flagella controls bacterial surface behavior"

**This includes:**

Materials and Methods
Supplemental Figures S1 to S8
Supplemental Tables S1 to S3

### Materials and Methods

#### Bacterial strains, plasmids, and growth conditions.

Bacterial strains used in this study are described in Table 1, the primers used are described in Table 2, and the plasmids used are described in Table 3. *P. aeruginosa* strains were grown on LB agar plates (1.5% Bacto Agar) and in liquid LB in a roller drum at 37°C. LB medium was prepared using premix Miller LB Broth (BD Biosciences) and using standard LB preparation protocols.

**Generation of *P. aeruginosa* mutants.** Gene deletions were generated using the lambda Red recombinase system as previously described (26). The deletion construct was Gibson-assembled from three PCR products. First fragment, approximately 500 bp upstream of the target insertion site was amplified from PA14 genomic DNA. Second fragment, containing *aacC1* ORF flanked by FRT sites was amplified from pAS03D. Third fragment, approximately 500 bp downstream of the target insertion site was amplified from PA14 genomic DNA. The Gibson- assembled product was transformed into PA14 cells expressing the plasmid pUCP18-RedS. The colonies were selected on 30 µg/ml gentamicin, the mutants of interest were counter-selected on 5% sucrose, and pFLP2 was used to flip out the antibiotic resistance gene. pUCP18-RedS and pFLP2 were selected for using 300 µg/ml carbenicillin.

**Construction of microfluidic devices.** Microfluidic devices were created using soft lithography techniques as previously described (26). Devices were designed on Illustrator (Adobe Creative Suite) and masks were printed by CAD/Art Services. Molds were made on 100 mm silicon wafers (University Wafer) and spin coated with SU-8 3050 photoresist (MicroChem). Polydimethylsiloxane (PDMS) chips were plasma-treated to bond with glass slides for at least 24 hours prior to experiments. The devices used in all experiments contained 7 parallel channels (500 µm width x 50 µm height x 2 cm length). Channels individually contained an inlet tube and an outlet tube. PDMS chips were plasma treated to a 60 mm x 35 mm x 0.16 mm superslip micro cover glass (Ted Pella, Inc.).

**Residence time assays.** Surface residence time experiments were performed as previously described (26). *P. aeruginosa* cells were loaded into plastic 5 mL syringes (BD) at mid-log phase with an optical density of approximately 0.5. All microfluidic experiments were performed at ~ 22 °C. The device set-up involves the loaded syringes attached to tubing which connects the needle to the inlet of the device (BD Intramedic Polyethylene Tubing; 0.38 mm inside diameter, 1.09 mm outside diameter). These syringes were situated on a syringe pump (KD Scientific Legato 210) which was used to produce fluid flow. The outlet of the device employed the same tubing and vacated into a bleach-containing waste container. The syringe pump was used to generate flow rates of 10 µl/min, which correspond to shear rates of 800 $s^{-1}$.

**Phase contrast microscopy.** Images were obtained with a Nikon Ti2-E microscope controlled by NIS Elements as previously described (26). All images were taken with Nikon 100x Plan Apo Ph3 1.45 NA objective, a Hamamatsu Orca-105 Flash4.0LT+ camera, and Lumencor Sola Light Engine LED light source.

**Imaging of flagella.** Strains were grown at 37°C overnight, back diluted 1:100, grown to early log phase and then incubated with Alexa488-mal (VWR) for 45 mins. Cells were washed twice using centrifugation to remove excess dye. *P. aeruginosa* cells were loaded into plastic 5 mL syringes (BD) at mid-log phase with an optical density of approximately 0.5. All microfluidic experiments were performed at ~ 22 °C. The device set-up involves the loaded syringes attached to tubing, connecting the needle to the inlet of the device (BD Intramedic Polyethylene

Tubing; 0.38 mm inside diameter, 1.09 mm outside diameter). Syringes were situated on a syringe pump (KD Scientific Legato 210) which was used to produce fluid flow. The outlet of the device employed the same tubing and vacated into a bleach-containing waste container. The syringe pump was used to generate flow rates of 10 µL/min, which correspond to shear rates of 800 $s^{-1}$. To view flagella, we employed a Nikon 100x/1.45 objective on an inverted fluorescent Nikon Eclipse Ti2 microscope with an ORCA-fusion BT sCMOS camera, for image acquisition. Flagellar images were taken using Nikon Elements Software and analyzed using ImageJ.

**Swim assay.** Strains were grown at 37°C overnight and then 1 mL aliquots were transferred to 1.5 mL microcentrifuge tubes. Sterile toothpicks were used to stab into LB plates containing 0.3% agar, and plates were incubated overnight at 37°C. The diameter of the swim zones was measured using Adobe Illustrator.

**Shear rate and shear force calculations.** The shear rate experienced in the microfluidic devices was calculated using the following equation:

$$Wall\ shear\ rate \sim \frac{6Q}{wh^2}$$

Where Q is the flow rate, w is the channel width, and h is the channel height. Shear stress was calculated as the product of shear rate and viscosity. Shear force was calculated as the product of shear stress and the surface area of a cell, which was estimated to be 2.5 $\mu m^2$ (9).

**Shear force modification assay.** To adjust the viscosity of *P. aeruginosa* cell cultures, we prepared a solution containing a 15% concentration of the viscous agent Ficoll. All microfluidic experiments were performed at ~ 22 °C. The device set-up involves the loaded syringes attached to tubing which connects the needle to the inlet of the microfluidic device (BD Intramedic Polyethylene Tubing; 0.38 mm inside diameter, 1.09 mm outside diameter). The loaded syringes were situated on a syringe pump (KD Scientific Legato 210) which was used to produce fluid flow. The outlet of the device employed the same tubing and vacated into a bleach-containing waste container. The syringe pump was used to generate flow rates of 1 µl/min (shear rate of 80 $s^{-1}$) and 10 µl/min (shear rate of 800 $s^{-1}$).

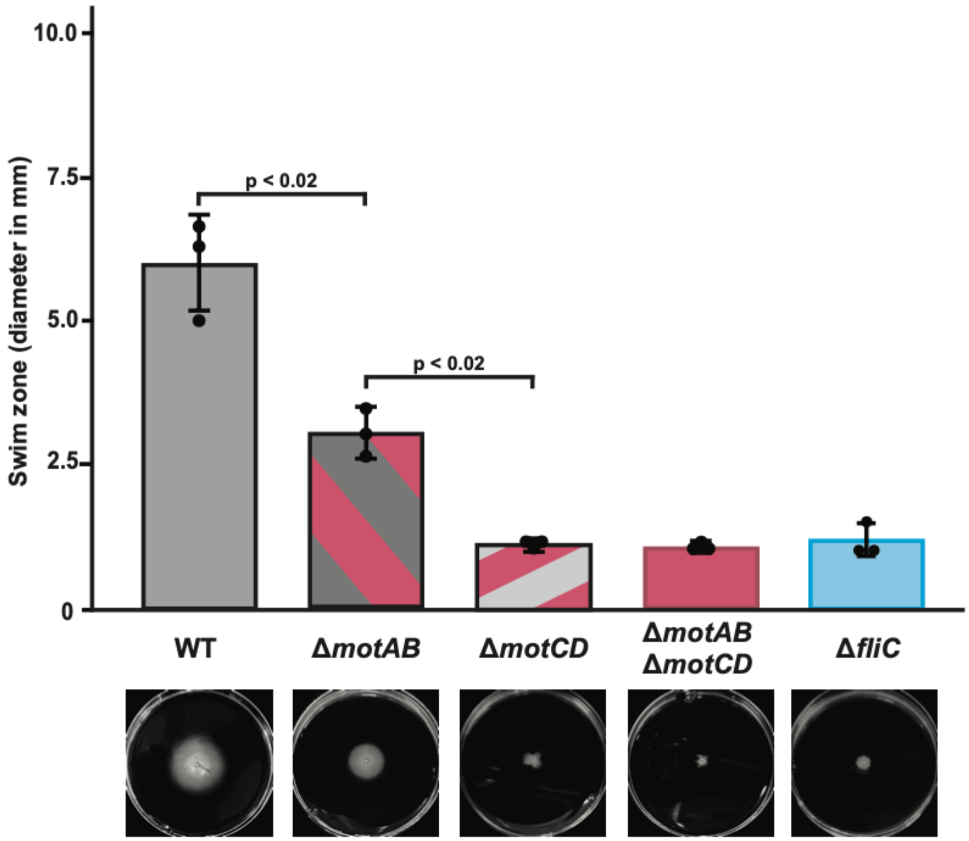


**Figure S1: MotAB and MotCD both contribute to swimming in agar motility assays.** Quantification of swim zones for WT and mutant strains. The diameter of each resulting swim zones was measured after 18 hours at 37°C. Quantification shows the average and SD of three biological replicates. P-values were calculated with the Student's T-test. Representative swim plate images for each strain were captured after 18 hours.

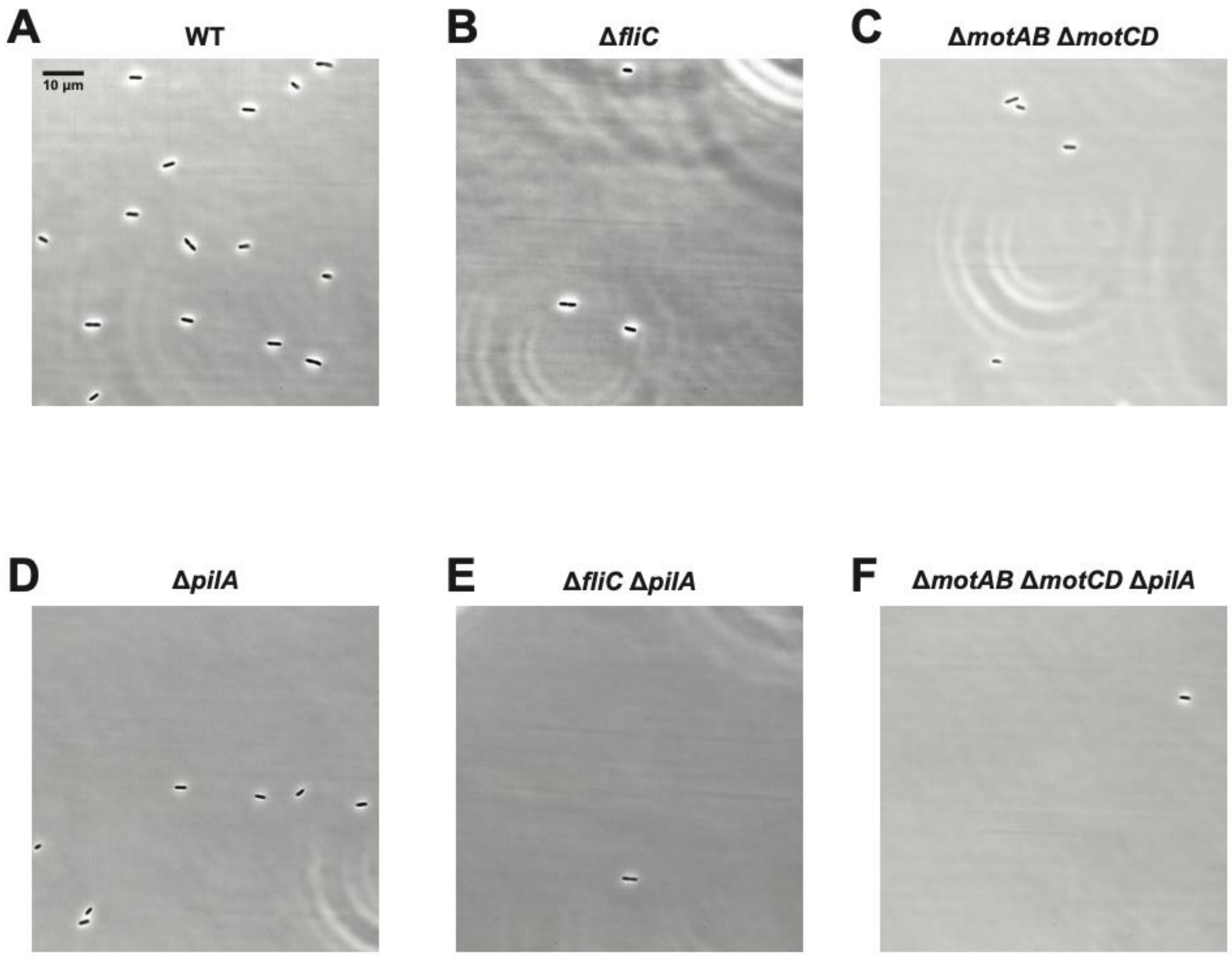


**Figure S2: Flagellar rotation promotes *P. aeruginosa* surface arrival in flow.** Representative phase images showing cells after they have arrived on the glass surface of the microfluidic device. Cells were flowed into the channel at a shear rate of 800 $s^{-1}$. Images were taken after 30 seconds of flow. Scale bar, 10 µm. Quantification of these experiments is shown in Figure 1B.

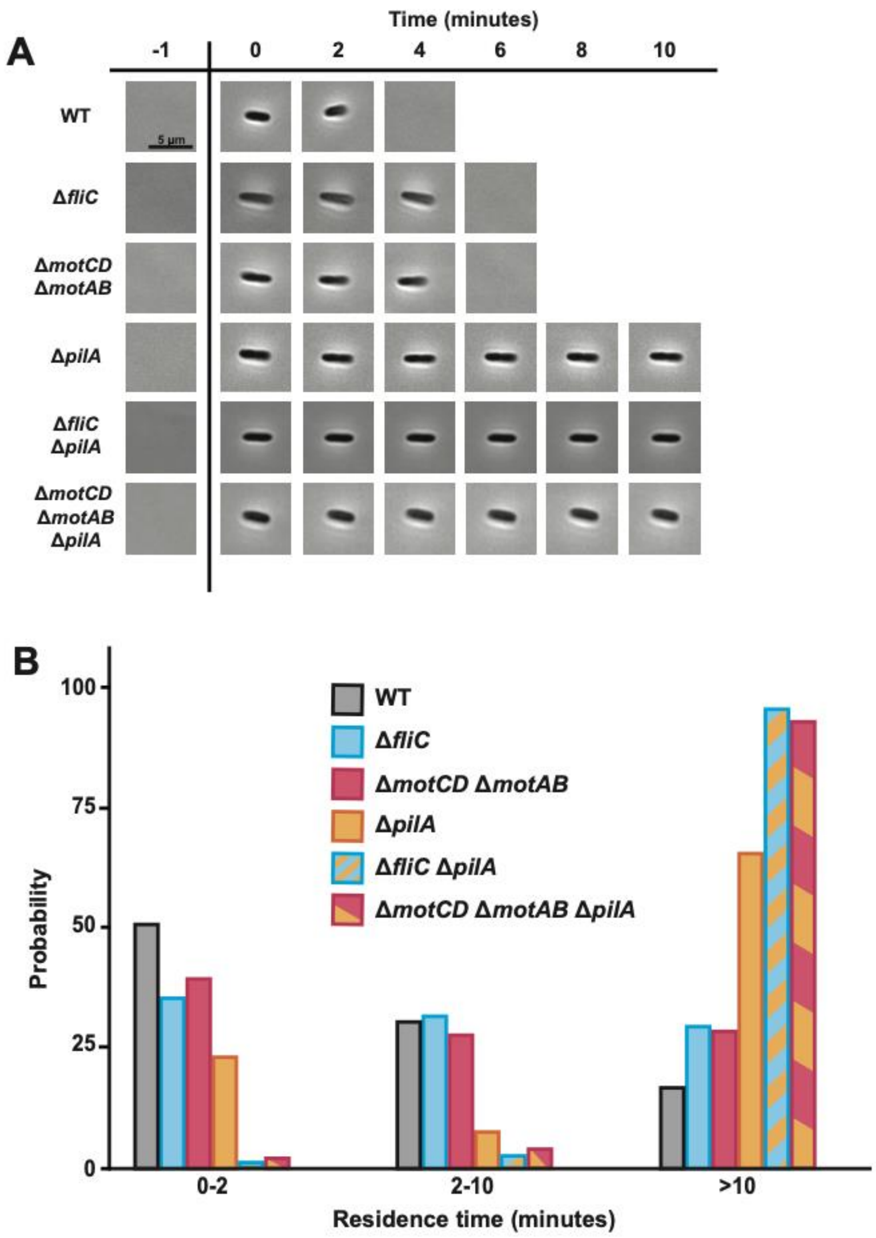


**Figure S3: Flagellar rotation promotes *P. aeruginosa* surface departure in flow.**
(A) Representative phase images of WT and mutant cells that arrived on the surface after being flowed into microfluidic devices at a shear rate of 800 $s^{-1}$. Scale bar, 5 µm. (B) Probability of the WT and mutant cell surface residence times from Figure 2 categorized into three groups (0-2 minutes, 2-10 minutes, and >10 minutes). Three biological replicates were performed and 150 cells (50 from each replicate) of each bacterial strain were chosen at random for quantification.

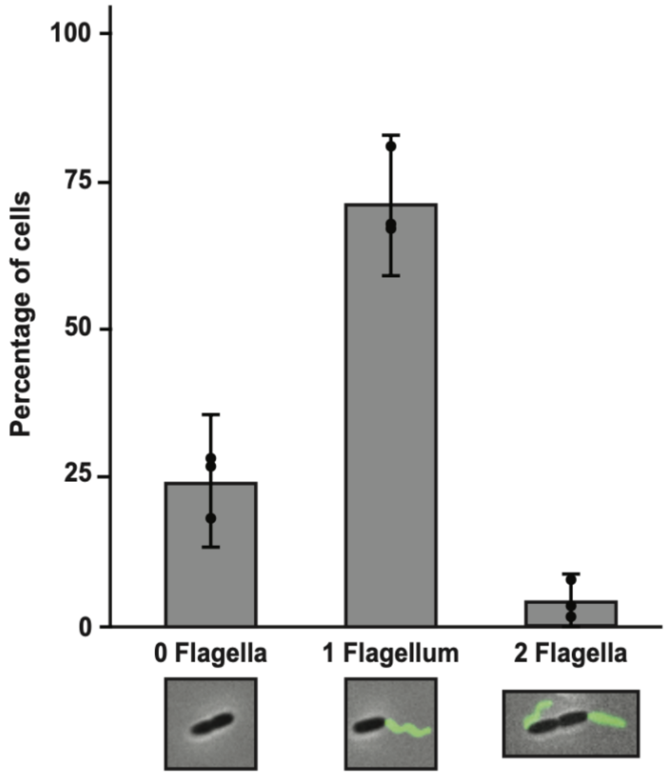


**Figure S4: WT *P. aeruginosa* cells typically have 1 flagellum.**
Quantification and representative images of surface-attached WT cells of three categories (no labeled flagella, one labeled flagellum, or 2 labeled flagella). Quantification shows the average and standard deviation of three biological replicates.

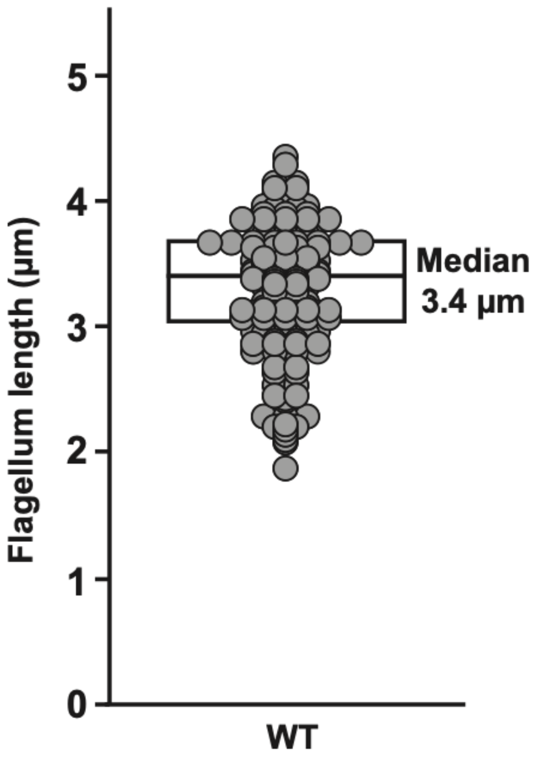


**Figure S5: Flagella of WT *P. aeruginosa* cells are typically 3.4 µm long.**
Quantification of flagellar length of fluorescently labeled WT cells. Flagellar length was measured from base to tip. Each data point represents the length of one flagellum. Three biological replicates were performed and 200 cells were chosen at random for quantification. The boxplot represents the 25$^{th}$ percentile, median, and 75$^{th}$ percentile.

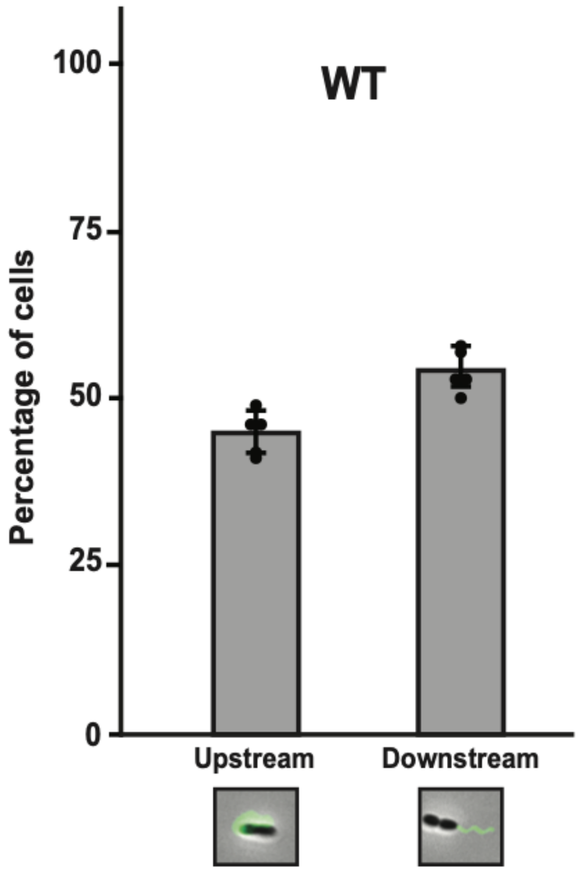


**Figure S6: Cells are equally likely to arrive with upstream or downstream facing flagella.** Quantification and representative images of surface-attached WT cells with upstream facing flagella or downstream facing flagella. Quantification shows the average and standard deviation of five biological replicates.

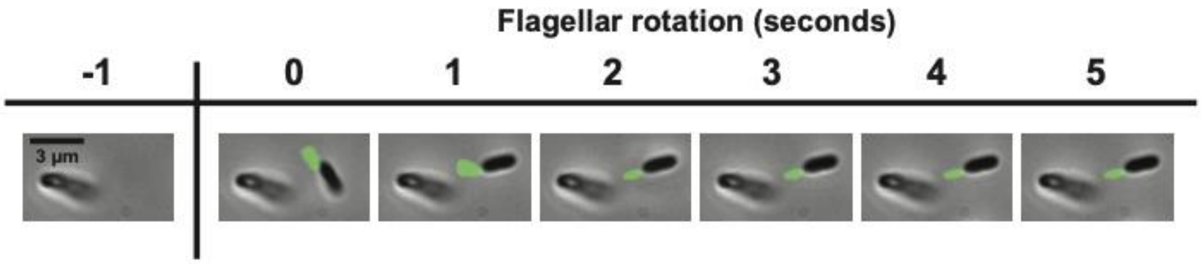


**Figure S7: Cell-cell shielding can prevent flow-induced flagellar bending.**
Images of a surface-attached WT cell with an upstream facing flagellum. Cell was flowed in at a shear rate of 800 $s^{-1}$. Cell arrives on the surface at 0 seconds and continues to rotate its flagellum for at least the next 5 seconds. Scale bar, 3 µm.

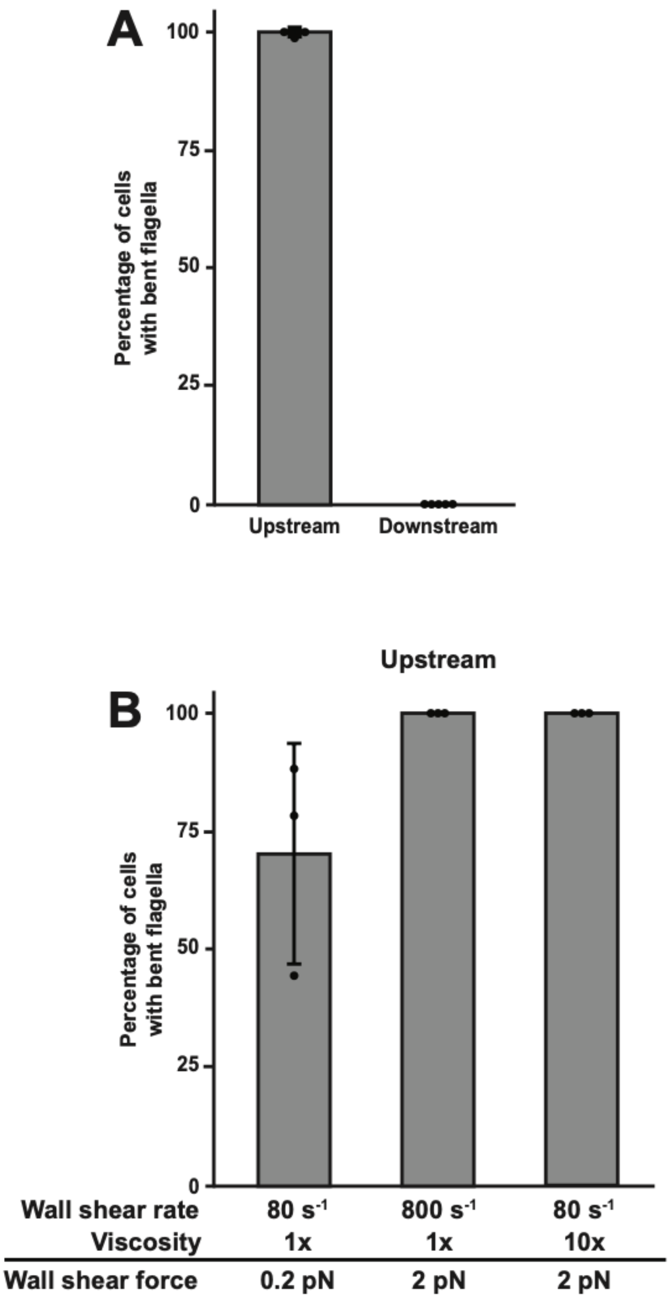


**Figure S8: Shear force bends upstream facing flagella around the cell.**
(A) Percentage of WT cells with bent flagella. Cells were flowed in at a shear rate of 800 $s^{-1}$ and categorized as having upstream facing or downstream facing flagellum. Quantification shows the average and SD of five biological replicates. (B) Percentage of upstream facing flagella that bent around cells during exposure to different shear forces. Shear force was increased either by changing shear rate or fluid viscosity. Shear rate was modified by changing the flow rate of the syringe pump. 10x viscosity was generated by adding 15% Ficoll, which has been shown previously to modify local viscosity (9, 42). Quantification shows the average and SD of three biological replicates.

**Table 1: *P. aeruginosa* strains used in this study**

| ***P. aeruginosa* strain** | **Description** | **Source** |
|---|---|---|
| PA14 | wildtype; clinical isolate from burn wound | (46) |
| JS177 | *ΔpilA::aacC1* | (26) |
| JS176 | *ΔpilA::FRT* | (26) |
| JS191 | *ΔfliC::aacC1* | This paper |
| JS190 | *ΔfliC::FRT* | This paper |
| JS197 | *ΔmotCD::aacC1* | This paper |
| JS200 | *ΔmotCD::FRT* | This paper |
| JS175 | *ΔmotAB::aacC1* | This paper |
| JS174 | *ΔmotAB::FRT* | This paper |
| JS206 | *ΔfliC::FRT ΔpilA::aacC1* | This paper |
| JS212 | *ΔfliC::FRT ΔpilA::FRT* | This paper |
| JS205 | *ΔmotCD::FRT ΔmotAB::aacC1* | This paper |
| JS213 | *ΔmotCD::FRT ΔmotAB::FRT* | This paper |
| JS318 | *ΔmotCD::FRT ΔmotAB::FRT ΔpilA::aacC1* | This paper |
| JS319 | Δ*motCD::FRT* Δ*motAB::FRT* Δ*pilA::FRT* | This paper |
| JS115 | *fliC* T394C; Cysteine knock-in mutant for flagellar labeling | This paper |
| JS327 | *fliC* T394C Δ*motCD::FRT* Δ*motAB::aacC1* | This paper |
| JS328 | *fliC* T394C Δ*motCD::FRT* Δ*motAB::FRT* | This paper |

**Table 2: Plasmids used in this study**

| **Plasmid** | **Description** | **Source** |
|---|---|---|
| pAS03D | Plasmid to generate deletion mutants in *P. aeruginosa* PA14 | (47) |
| pFLP2 | Plasmid expressing FLP2 to recombine FRT sites | (48) |
| pUCP18-RedS | Lambda red recombineering vector | (47) |

**Table 3: Primers used in this study**

| Primer | Sequence (5'-3') |
|---|---|
| Δ*pilA*-1 | cgcagtaggcgataccgaat |
| Δ*pilA*-6 | aggaactcggttttctccgc |
| Δ*fliC*-1 | ctgctatcgcgacagtctcc |
| Δ*fliC*-6 | aatcggtcgagcctactcct |
| Δ*motCD*-1 | ggtgctgatccagcacatgcc |
| Δ*motCD*-2 | ccatcgaaggcagtctcctcattcc |
| Δ*motCD*-3 | **gcaggaatgaggagactgccttcgatggccacccacgtga**attccggggatccgtcgacc |
| Δ*motCD*-4 | **tggccagttcggccccggcgcccgtacgcaaaccatgttt**gtgtaggctggagctgcttc |
| Δ*motCD*-5 | gcgcgaaacatggtttgcgtac |
| Δ*motCD*-6 | cattgaccatcagcacgccgag |
| Δ*motAB*-1 | cagtggatttcctgccagagc |
| Δ*motAB*-2 | gaggaccggacgtgcgaaatg |
| Δ*motAB*-3 | **tccccctccttatcctgttcatttcgcacgtccggtcctc**attccggggatccgtcgacc |
| Δ*motAB*-4 | **gcacattctggcaagcatgccggaaacggactactgaacc**gtgtaggctggagctgcttc |
| Δ*motAB*-5 | ggttcagtagtccgtttccggc |
| Δ*motAB*-6 | catcgatgcgctgctgaatgt |
